# Protective efficacy of calcium phosphate nanoparticle adsorbed bivalent subunit vaccine of *Pasteurella multocida* against homologous challenge in mice

**DOI:** 10.1101/2020.09.06.284687

**Authors:** Songyukta Shyam, Shantanu Tamuly, Probodh Borah, Rajeev Kumar Sharma

**Affiliations:** Department of Animal Biotechnology, College of Veterinary Science, Assam Agricultural University, Khanapara, Guwahati. 781022 (INDIA); Department of Veterinary Biochemistry, College of Veterinary Science, Assam Agricultural University, Khanapara, Guwahati. 781022 (INDIA); Department of Veterinary Microbiology, College of Veterinary Science, Assam Agricultural University, Khanapara, Guwahati. 781022 (INDIA)

**Keywords:** calcium phosphate nanoparticle, *Pasteurella multocida*, outer membrane proteins, swine pasteurellosis, vaccine

## Abstract

Swine pasteurellosis, caused by *Pasteurella multocida* capsular types A and D, causes heavy economic loss to the pig farmers. The vaccine presently used is abacterin of *Pasteurella multocida* capsular type B that is proven to be effective against bovine pasteurellosis. However, its efficacy against swine pasteurellosis is questionable. The present study was carried out to evaluate the efficacy of calcium phosphate nanoparticle adjuvanted bivalent subunit vaccine prepared from *Pasteurella multocida* capsular types A and D along with a monovalent subunit vaccine prepared from *Pasteurella multocida* capsular type B in mice. The Alum precipitated bacterin vaccine was used as the control. The bivalent subunit vaccine comprising the immune components of both the capsular types showed significantly higher IgG response than either of the other two vaccines. Both the calcium phosphate nanoparticle adjuvanted vaccines could elicit 100% protection in mice against homologous challenges but the aluminum hydroxide adjuvanted bacterin vaccine could not elicit significant protection. Based on this preliminary work, it was concluded that the bivalent subunit vaccine would be a better option for immunization of swine against swine pasteurellosis.

**IMPORTANCE OF THE WORK:** The swine pasteurellosis is an important economic disease affecting the pig population in the North-eastern part of India that contributes the major pig population. The disease is caused by Serotype A and D of *Pasteurella multocida*. At present the inactivated vaccine is used that is actually developed against P_52_ strain of serotype B:2 of *Pasteurella multocida*, which is mainly involved in haemorrhagic septicaemia (or bovine pasteurellosis) that affects the cattle, buffaloe, sheep and goat. As a result, the present vaccine does not give sufficient protection in pigs but gives significant protection in cattle, buffaloe, sheep and goat. Hence, there is a need of development of vaccine that can address specifically swine pasteurellosis by targeting serotype A and D of *Pasteurella multocida*.

## INTRODUCTION

Swine pasteurellosis is caused by *Pasteurella multocida* capsular types A and D,while the capsular type B:2 of *P. multocida* is the causative agent of haemorrhagic septicaemia (HS) that affects mainly cattle, buffalo, sheep and goat. Swine pasteurellosis is responsible for significant economic loss to the pig industry in the north-eastern region of India. The disease occurs in acute septicaemic as well as subacute pneumonic forms. Pneumonic forms are caused by *P. multocida* of capsular types A and D^1 cited in 2^. There are also reports of acute septicaemia caused by capsular type B organism^3cited in 2^. Swine pasteurellosis often occurs as complications with swine fever, swine influenza and may also be associated with parasitic, viral and other bacterial diseases or environmental factors that impair pulmonary function.

Presently in India, an alum- or oil-adjuvanted killed vaccine prepared from *P. multocida* serotype B: 2 (P_52_ strain) is used for control of swine pasteurellosis in endemic areas. This vaccine is unable to elicit sufficient protective immune response against swine Pasteurellosis as it is prepared from P_52_ strain of serotype B: 2 of *P. multocida* that is efficient in controlling bovine pasteurellosis. These vaccines have certain limitations, *viz*. alum precipitated vaccine induces local reactions, elicits IgE antibody responses and generally fail to induce cell mediated immunity. Therefore, the duration of immunity induced is only for 4 to 6 months. Alum type adjuvants are not effective for all antigens. Whereas, oil adjuvanted vaccines being too viscous are difficult to inject in animals particularly during herd vaccination and it has site-specific and pyrogenic responses. Outbreaks are still occurring in endemic areas even after vaccination. The commercial vaccines prepared from *P. multocida* serotype B: 2 (P_52_ strain) has been successful in controlling HS but this serotype is generally not involved in swine pasteurellosis. This necessitates the exploration of other vaccine candidates that may ensure adequate protection against *P. multocida* capsular types A and D.

Live vaccines have also been used in some areas but outbreaks attributed to vaccine strains and sometime reversal to the virulent form limit the use of these vaccines. Live vaccines using capsular type B: 3, 4 strains have been reported to be unsafe for primary vaccination in young calves. The inactivated form of vaccine is widely used throughout the world. Generally, they are poor immunogens thus require efficient adjuvant system. Therefore, the development of more efficient and safe adjuvants, and vaccine delivery systems to obtain high and long-lasting immune response is of utmost importance.

Various workers aimed at developing more effective vaccines by identifying potential protective antigen(s) and formulating new vaccine adjuvants. The capsular polysaccharide has shown induction of higher level of protection in rabbits when conjugated with bovine serum albumin. Lipopolysaccharide of *P. multocida* has been found to be partially immunogenic in mammals. Cattle and rabbits have not been readily protected against *P. multocida* infection following immunization with LPS. Protein is an integral part of the outer membrane of gram-negative bacteria and is found to be associated with immunity^4, 5, 6^. Many different studies on outer membrane proteins (Omp) have shown that they are major immunogens against homologous challenges in mammal^7^. The omps being non-living vaccine candidates require an efficient adjuvant system for eliciting efficient immune response^8^.

An ample amount of work has been carried out on oil and alum adjuvants, which have respective drawbacks of poor syringability and site-specific reactions^8^. Some works were carried out on exploration of calcium phosphate nanoparticles as an adjuvant. Nanoparticles have recently been shown to possess significant potential as a drug delivery system. Studies have showed significant adjuvanticity of calcium phosphate nanoparticles when complexed with whole capsid proteins of virus and recombinant protein of bacteria^9, 10, 11^. Calcium phosphate is a natural component the body and known to act by depot effect, anddue to its smaller size, facilitates efficient uptake by macrophages. The potential advantages of calcium phosphate nanoparticles as an adjuvant are-1. Reduce systemic side effect, 2. Efficiently deliver antigens to antigen presenting cells (APCs), especially dendritic cells 3. Provide sustained release of the encapsulated antigens (depot effect)^12^.

In the present study, the whole outer membrane proteins obtained from capsular types A and D (isolated from upper respiratory tract of pig)as well as the bacterin prepared from P_52_ vaccine strain of *P. multocida* were conjugated with calcium phosphate nanoparticles and their protective efficacy was evaluated in mice.

## MATERIALS AND METHOD

### Bioethics permission

The study was approved by the Institutional Animal Ethics Committee of Assam Agricultural University, Khanapara, Guwahati-781022 (India) vide approval No. 770/ac/CPCSEA/FVSc/AAU/IAEC/13-14/191 dated 10-2-2014.

### Bacterial strains and media

The bacterial strains used in the present study were P_52_ strain (capsular type B:2) of *P. multocida*, swine isolates of capsular types A and D of *P. multocida* obtained from the repository of Department of Veterinary Microbiology, College of Veterinary Science, Assam Agricultural University, Guwahati. The bacterial isolates were grown in brain heart infusion broth (Hi-Media) at 37°C in shaking condition to mid-logarithmic phase.

### Isolation of whole outer membrane protein

The whole outer membrane proteins were purified from revived isolates of capsular types A, B, and D of *P. multocida*. Briefly, the glycerol stock of the isolates were streaked onto a blood agar plate and incubated overnight at 37°C. The single colonies were inoculated into brain heart infusion broth (Hi-Media) and incubated at 37°C in shaking condition till the mid-log phase. Then 0.1 ml of culture was inoculated into Swiss Albino mice (Lacca strain) intraperitoneally. The organism was re-isolated from the heart blood, and its species and capsular type was confirmed by PM specific PCR^13^ and capsular type specific PCR,^29^ respectively. The whole outer membrane proteins were extracted from all the three strains as per the method described by Choi-Kim *et al*.^15^. The protein was quantified by the method described by Lowry^16^. The presence of proteins were documented by one dimensional SDS-PAGE, using 5% stacking gel and 12% separating gel^17^.

### Preparation of Calcium Phosphate Nanoparticle-adsorbed Outer Membrane Protein Conjugate (CAP-Omp)

One mg of outer membrane protein (0.5 mg of OMP each from *P. multocida* of capsular types A and D) was lyophilized in sterile conical flasks. This was followed by addition of 7.5 ml of 12.5 mM calcium chloride, 7.5 ml of 12.5 mM dibasic sodium phosphate solution and 1.5 ml of 15.6 mM sodium citrate. Flask containing the suspension was stirred till the average particle size was less than 100 nm. The suspension was centrifuged at 800×g for 20 minutes at 4°C. The supernatant was discarded and the pellet was washed with 0.1M PBS (pH 7.2). One ml cellubiose (129 mM) was added to the pellet and kept for overnight at 4°C. Four mg of OMP was added to the pellet with cellubiose and was incubated at 4°C for 1 hour. The suspension was centrifuged at 800×g for 20 minutes and the pellet was lyophilized and kept in −20°C till further use. The calcium phosphate nanoparticle-OMP conjugate containing the OMP of P_52_ strain was made in a similar manner.

### Characterization of Nanoparticles

The characterization of nanoparticles was done by transmission electron microscopy in Indian Institute Technology, Guwahati. For transmission electron microscopy (TEM-2100F), one drop of calcium phosphate nanoparticle suspension was put on parafilm. The 200 mesh copper carbon grid was soaked into the drop. The grid was placed on the filter paper and the grid was allowed to dry for one hour. The dried grid was then placed inside the electron microscope chamber after creation of vacuum. The particles were viewed at 150,000 Xmagnification.

### Estimation of Protein Load in the Complex (CAP-Protein)

The amount of protein load of calcium phosphate nanoparticle-OMP complex was measured by using protocol of Joyappa *et al*.^18^. The known amount of calcium phosphate nanoparticle-OMP complex pellet was taken in a microcentrifuge tube. The pellet was suspended in 100mM EDTA solution and incubated at room temperature for 30 minutes. The suspension was centrifuged at 800×g for 20 minutes. The supernatant was taken in a fresh tube and protein was estimated at 280 nm by spectrophotometry.

### Immunization Protocol in mice

Twenty four Swiss Albino mice were divided randomly into four groups containing six mice in each group as depicted in Table 1.

**Table 1.** Groups of mice injected with different vaccine formulations and their route of inoculation

| Groups | Material injected | Route |
| --- | --- | --- |
| Group A | Calcium phosphate nanoparticle-outer membrane protein (capsular type A and D) complex (CAP-OMP(A+D)) | s/c |
| Group B | Calcium phosphate nanoparticle-outer membrane protein complex of P<sub>52</sub> strain (CAP-OMP(P<sub>52</sub>)) | s/c |
| Group C | Calcium phosphate nanoparticle ( without incorporation of protein) (CAP-Void) | s/c |
| Group D | Alum precipitated bacterin vaccine. | i/m |

The vaccine formulations were normalized to 1μg per ml. The mice belonging to groups A and B were injected with 100 μl of vaccine preparation containing 100 μg of OMP of *P. multocida* of capsular types A and D, and 100 μg of OMP of serotype B: 2 (P_52_ strain), respectively. The mice in group C were injected with calcium phosphate nanoparticles (without incorporation of protein) dissolved in 0.1M PBS (pH7.2). The mice of group D were injected subcutaneously with 0.2 ml of commercial alum adjuvanted bacterin vaccine (obtained from Department of Veterinary Microbiology, College of Veterinary Science, Assam Agricultural University).

The booster dose was administered on 14^th^day after primary vaccination. Blood samples were collected from the tail vein of mice on the day of primary vaccination and on 7, 14, 21 and 28 days post primary vaccination. The serum was separated and stored at −20°C till further use.

#### Enzyme-Linked Immuno-Sorbant Assay

The serum IgG titre was tested indirect ELISA. Briefly, the ELISA 96 wells polysorb microtiter plate was coated with either 200 ng of OMP of *Pasteurella multocida* capsular type A or D or B (P_52_) or 32 μg of whole cell sonicated lysate of P_52_ strain of *Pasteurella multocida*. The coated plates were stored at 4°C for overnight.

The plate was washed thrice with washing solution (PBS containing 0.05% Tween-20 pH 7.4). Hundred μl of blocking solution (2% BSA in PBS) was added to each well and the plate was kept at 37°C for one hour. The plates were washed thrice with washing solution. Then 100 μl of serially diluted serum was added to each well and the plates were incubated for 2 hours at 37°C. The plates were washed thrice with washing solution. Hundred μl of anti-mouse HRP conjugated IgG antibody (1:2000) was added to each well and the plates were kept for incubation for one hour at 37°C. The OPD substrate solution (24.3 mM citric acid, 51.4 mM dibasic sodium phosphate, 0.4 mg per ml orthophenyl diamine, 0.4 μl per ml H_2_O_2_) was prepared during the time of incubation. The plates were washed thrice with washing solution. Then 100 μl of OPD substrate was added and the plates were incubated at room temperature in dark for 30 minutes. Then 100μl of 1.5N NaOH was added to each well. The OD was read in 492 nm. The cut-off value was determined by mean OD of negative control plus 2×Standard deviation. The titre was determined using regression analysis taking OD value as predictor variable (x-variable) and reciprocal of log_10_ of serum dilution as response variable (y-variable).

#### Challenge Studies in Mice

All groups were inoculated intra-peritoneally with virulent *P. multocida* capsular types A or D at the dose rate of 100×LD_50_ on 28^th^ day post primary vaccination. The animals were observed for mortality or any kind of clinical symptoms for 72 hours. The re-isolation of organism was attempted from the heart blood of dead mice using the method as mentioned above under the section “Isolation of whole outer membrane protein”.

#### Statistical Analysis

The difference in humoral immunity between the groups were analyzed by one way ANOVA and among the different days within the same group by one way repeated measures ANOVA. The analysis of results of the animal challenge study was performed by Chi-square test of independence using Yates’s correction. The p-value of less than 0.05 was considered significant and less than 0.01 was considered highly significant. The analysis was performed with the statistical software R (R Core Team, 2019).

## RESULTS

### Protein profile of outer membrane protein of *Pasteurella multocida* capsular types A, D and B

Protein profiles of OMPs were studied by employing 12.5% SDS-PAGE. Coomassie brilliant blue stained gel revealed six prominent OMP bands of capsular type A having the relative molecular weights (Mr) ranging from 28.88 to 63.7 kDa; seven prominent OMP bands of capsular type D having Mr ranging from23.88 to 94.76 kDa and three prominent OMP bands of serotype type B: 2 (P_52_ strain) having Mr ranging from20.38 to 47.95 kDa. One OMP of Mr 47.95 kDa was found to be commonly expressed in all the three capsular types. On the other hand, the OMP of molecular weight 28.88 kDa was found to be commonly expressed in capsular type A and P_52_ strain of *P. multocida*.

**Table. 2. :** Relative molecular weights of OMP bands of capsular type A, D and P_52_ strain of *Pasteurella multocida*

| Relative molecular weights of OMPs in Kilo-Daltons (kDa) on the basis of electrophoretic migration in SDS-PAGE |  |  |
| --- | --- | --- |
| Capsular type A | Capsular type D | P<sub>52</sub> vaccine strain |
| 63.7 | 94.76 | 47.95 |
| 55.29 | 75.91 | 33.83 |
| 47.95 | 54.42 | 20.38 |
| 41.57 | 47.95 |  |
| 33.83 | 42.24 |  |
| 28.88 | 28.88 |  |
|  | 23.88 |  |

**Fig. 1.**
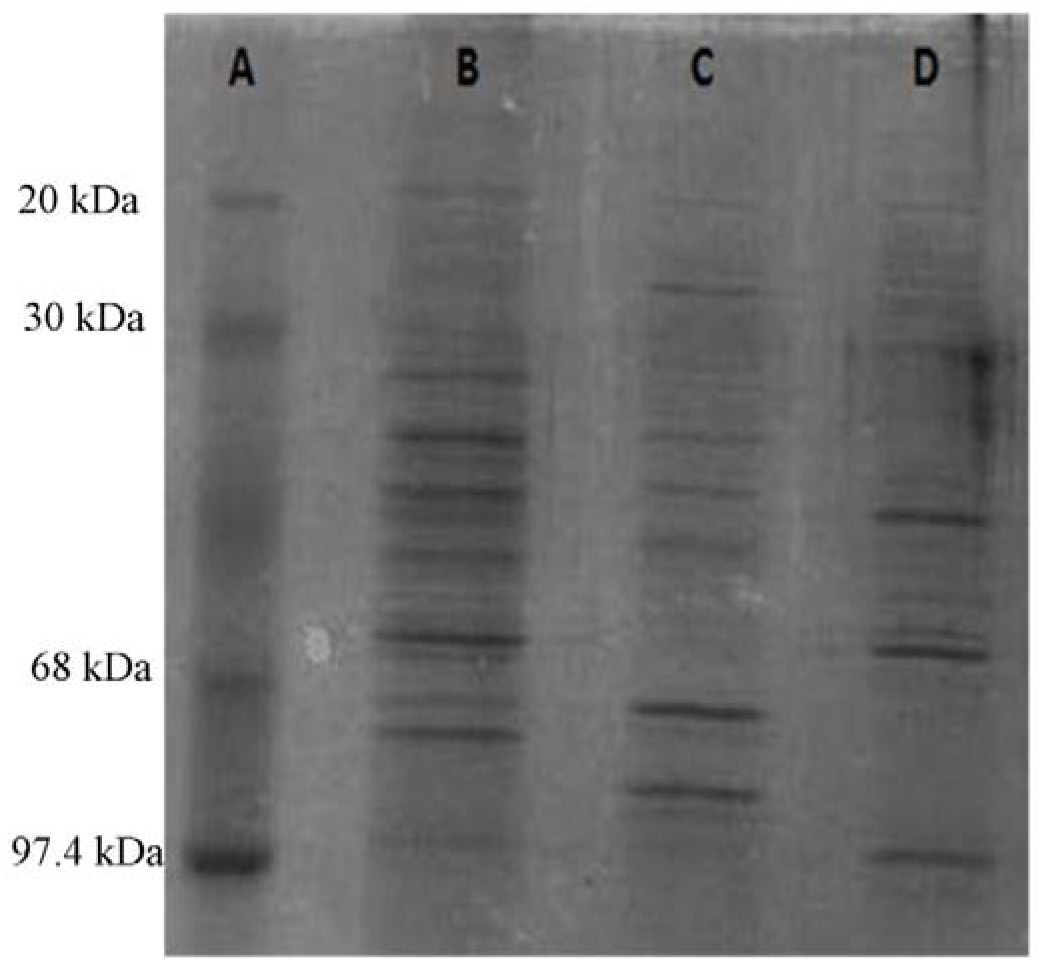
Coomassie Brilliant Blue stained SDS-PAGE (12.5%) profile of outer membrane proteins of *P. multocida* of different capsular types along with standard MW markers (Lane A =molecular marker; Lane B=*P. multocida* capsular type A; Lane C= *P. multocida* capsular type D; Lane D= *P. multocida* capsular type B: 2)

### Production of *P. multocida* outer membrane protein loaded calcium phosphate nanoparticle

The conjugate of protein and calcium phosphate nanoparticle was characterized by transmission electron microscopy. Majority of nanoparticles were of spherical morphology with a diameter of 39.9 nm to 80 nm. The distribution of size of the nanoparticlesis depicted in Fig. 2. A total of 10 μg protein could be loaded in 1 mg of calcium phosphate nanoparticles.

**Fig. 2.**
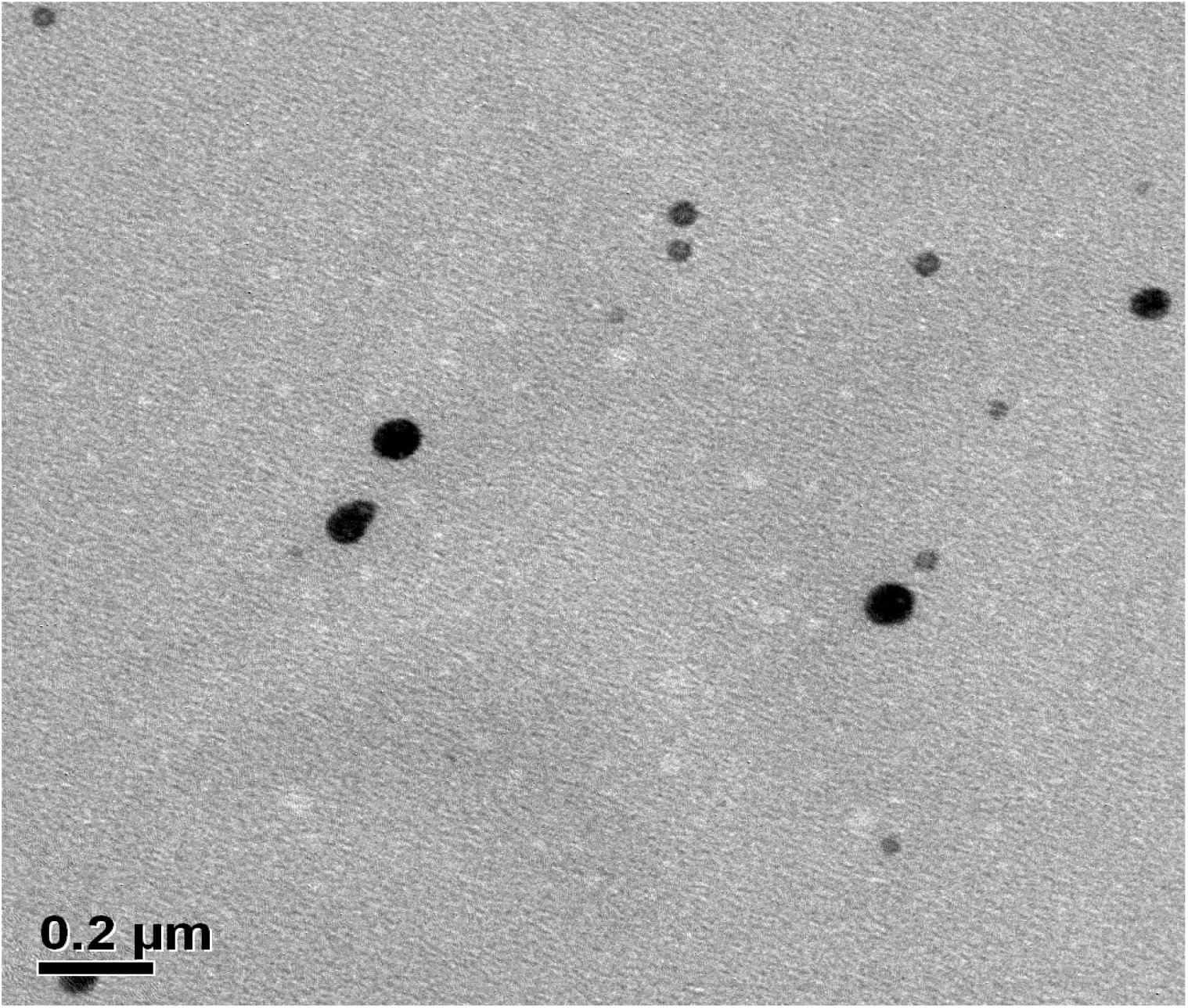
Electron micrograph of CAP-OMP complex at one hour stirring. The particle size was about 39.9nm to 80nm (150,000 × magnification).

### Induction of anti-IgG response in mice

The mean IgG titre against OMP of capsular type A in the group of mice vaccinated with CAP-OMP-(A+D) increased up to 14 days after the primary vaccination. Following booster vaccination on the 14^th^day, it increased slightly up to the 21^st^day which started declining on 28th day post primary vaccination; but there was no significant difference between the IgG titres on 21st day and 28th day (P > 0.05). The mean IgG titre against OMP of capsular type D in the same group increased up to 14 days and was maintained till 28 days as there were no significant difference between the titres recorded on 14^th^day and 21^st^ dayas well as between 21^st^day and 28^th^day (P >0.05).

The group of mice injected with CAP-OMP-(P_52_) showed similar pattern of IgG response as shown by CAP-OMP-(A+D) having the correlation of 97.94% and 98.62% for IgG response against OMP of CapA and against CapD, respectively. The mean IgG titre against OMP of capsular type A was found to be significantly higher in the group injected with CAP-OMP-(A+D) than that of the group injected with CAP-OMP-(P_52_) (P <0.01). The mean IgG titre of the group of mice injected with bacterin vaccine was found to be significantly lessercompared to that of the other two vaccinated groups.

The mean IgG response in all the groups against OMP of capsular type D increased up to 7 days post-primary vaccination and then it was maintained up to the 28^th^day. The mean IgG titre of the alum-adjuvanted bacterin vaccinated group rose up to 14 days but it started declining from 21 days even after booster vaccination on the 14^th^day post-primary vaccination.

**Fig. 3.**
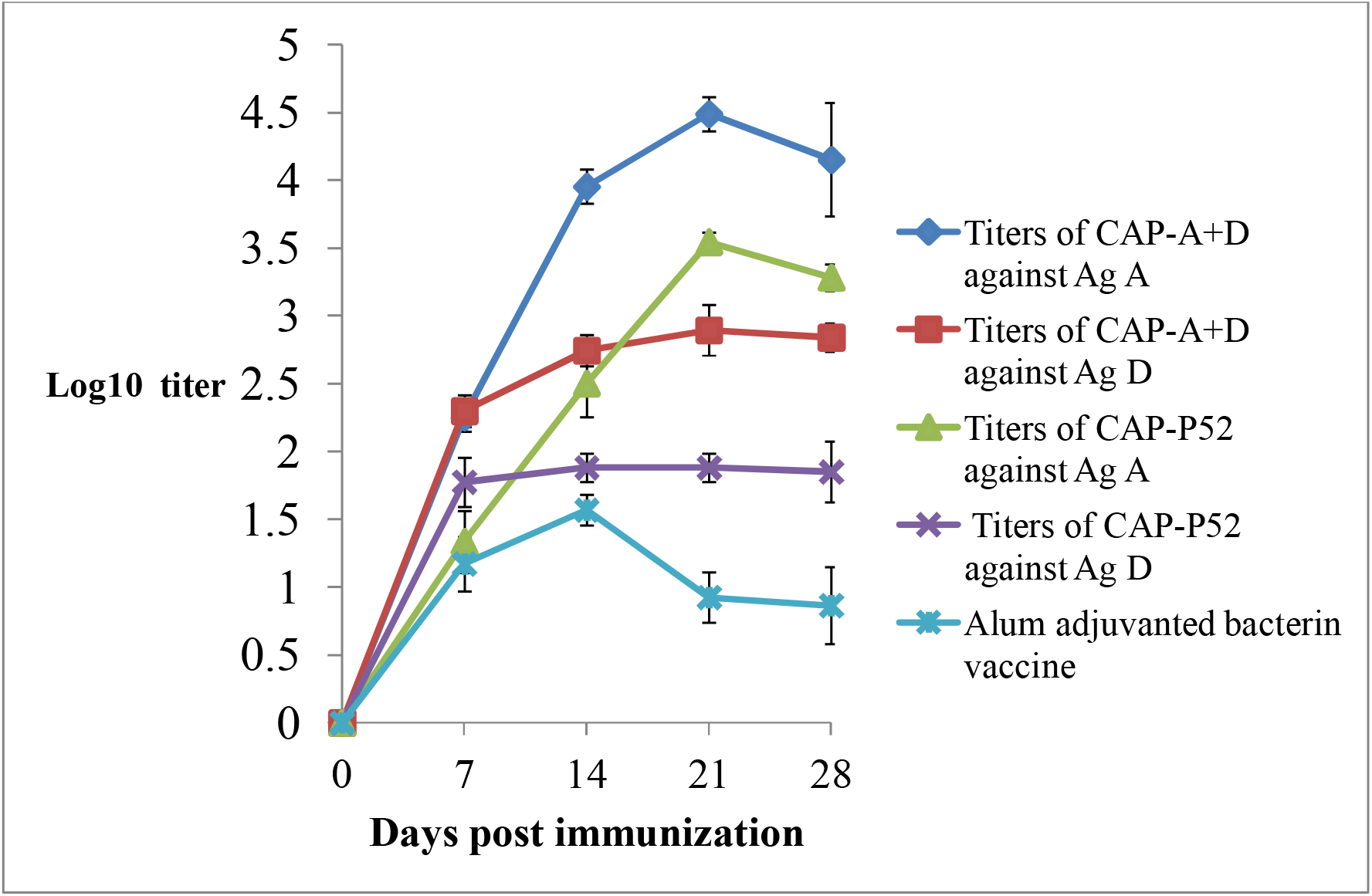
The graphical representation of mean log10IgG titre of different groups of mice against outer membrane protein of *Pasteurella multocida* capsular type A or D.

**Table 3.**
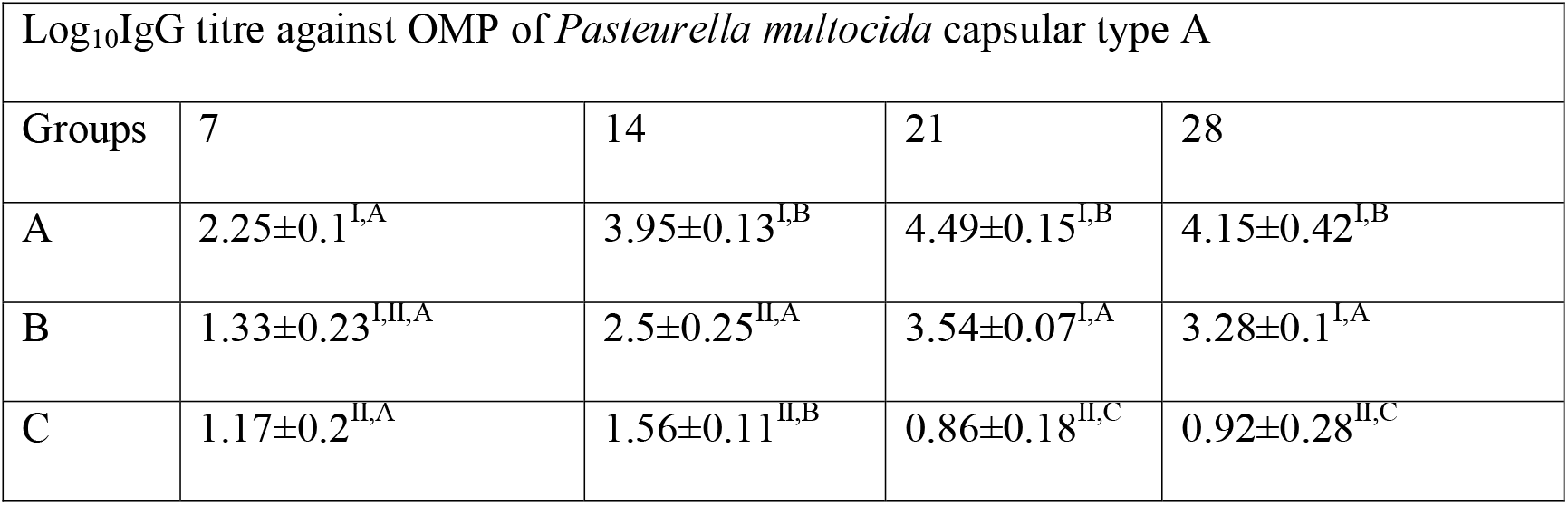
The tabular representation of mean log_10_ IgG titre of different groups of mice against outer membrane protein of *Pasteurella multocida* capsular type A (The different alphabets between column and different roman numbers between different rows indicate statistically significant difference (P <0.05)).

**Table 4.** The tabular representation of mean log_10_ IgG titre of different groups of mice against outer membrane protein of *Pasteurella multocida* capsular type D (The different alphabets between column and different roman numbers between different rows indicate statistically significant difference (P <0.05)).

| IgG titre Against OMP of *Pasteurella multocida* capsular type D |  |  |  |  |
| --- | --- | --- | --- | --- |
| Groups | 7 | 14 | 21 | 28 |
| A | 2.3±0.12<sup>I,A</sup> | 2.74±0.12<sup>I,A</sup> | 2.89±0.19<sup>I,A</sup> | 2.84±0.1<sup>I,A</sup> |
| B | 1.77±0.18<sup>III,A</sup> | 1.88±0.1<sup>III,A</sup> | 1.88±0.1<sup>III,A</sup> | 1.85±0.23<sup>II,A</sup> |
| C | 1.17±0.2<sup>II,III,A</sup> | 1.56±0.11<sup>II,III,A</sup> | 0.86±0.18<sup>II,A</sup> | 0.92±0.28<sup>III,A</sup> |

### Challenge study

The challenge study indicated that both the vaccine formulations containing calcium phosphate nanoparticles as adjuvant elicited 100% protection against challenge with either *P. multocida* capsular type A or D. On the other hand, the mice belonging to the group injected with alum adjuvanted bacterin showed the survivability of 50% and 33.33% when challenged with *P. multocidaof* capsular types A and D, respectively.

**Table 5.**
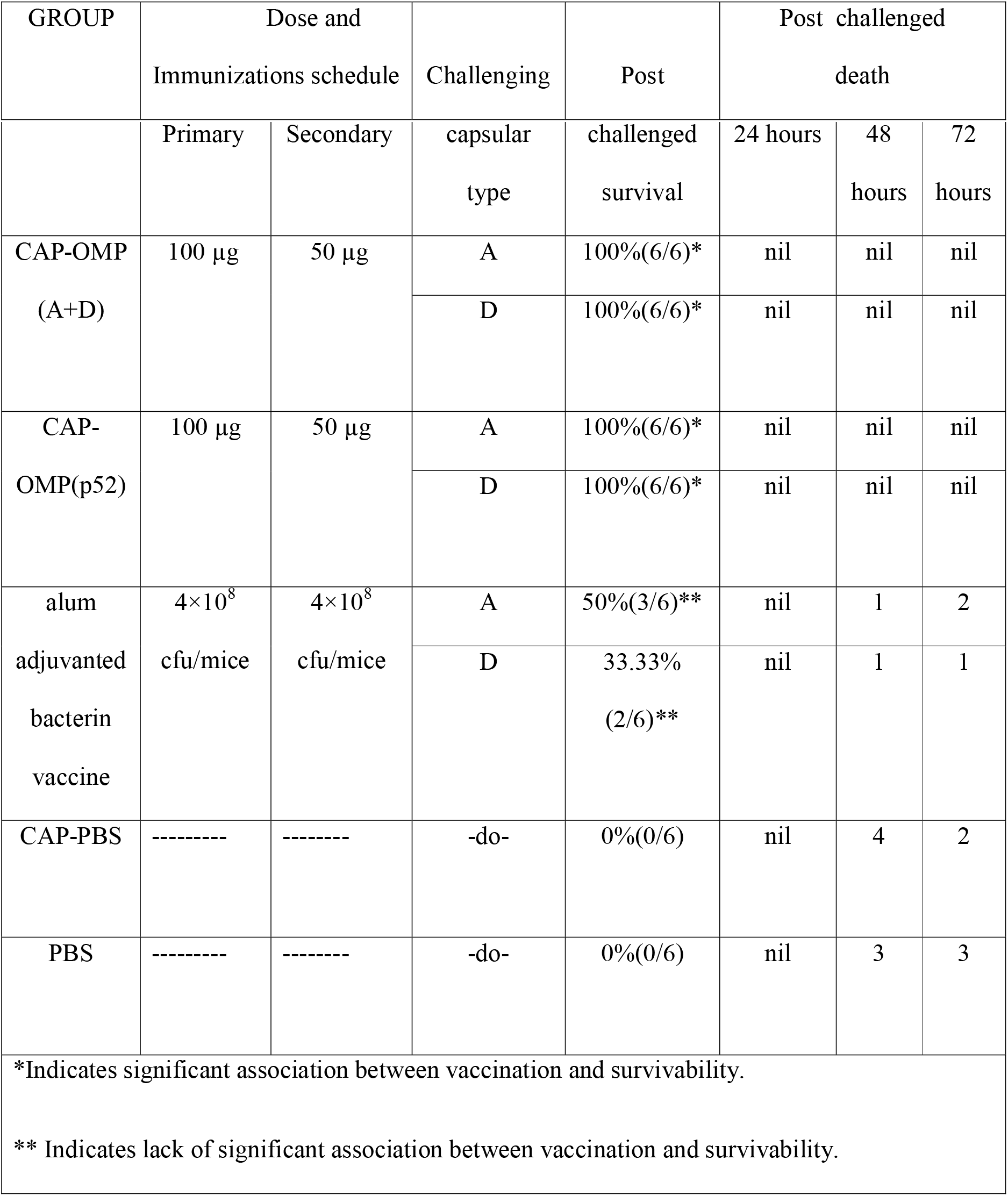
Protection of mice immunized with calcium phosphate nanoparticles -OMP of capsular types A and D and calcium phosphate nanoparticles - OMP of P52 and alum adjuvanted bacterin vaccine

## DISCUSSION

*Pasteurella multocida* of serotype B: 2 (P_52_ strains) is widely recognized as a causative agent of haemorrhagic septicemia of cattle and buffaloes^9^. Capsular types A and D cause economic losses in swine because of their association with progressive atrophic rhinitis and enzootic pneumonia^1^. Their association with acute septicaemia in pigs has been recognized^3^. *Pasteurella multocida* is a part of the commensal microflora in the upper respiratory tract of pigs. The organism was shown to appear intermittently in the naso-pharynx and subsequently shed in nasal secretions^20^. During such period, the carrier animal becomes a source of infection for in-contact susceptible animals. *Pasteurella multocida* has been classified into five serogroups on the basis of capsular antigens-A, B, D, E and F^21^. Besides their variable geographical distribution, these serogroups are more or less specific with regard to their host.^4^ Capsular types A & D cause economic losses in the swine industry because of their association with progressive enzootic pneumonia, also known as pneumonic pasteurellosis while atrophic rhinitis is associated with the capsular type D^23^.

Although several vaccines are used to control bovine pasteurellosis (Haemorrhagic septicemia), no vaccine is currently available for controlling swine pasteurellosis in India. Therefore occurrence of the disease in many regions of the country has been reported^18, 24, 25^. The SDS-PAGE banding patterns of OMP of *Pasteurella multocida* of capsular types A, D and B in the present study revealed significant differences which could be the possible reason behind the less efficiency of conventional vaccine prepared from capsular type B (P_52_ strain) in protecting pigs against swine pasteurellosis. As both the capsular types of A and D are involved in swine pasteurellosis, it would be prudent to use a bivalent vaccine containing the immune components of both the capsular types. The present study was carried out to assess the comparative efficacy of three vaccine formulations, *viz*. the calcium phosphate nanoparticle adjuvanted OMP based vaccines, either bivalent (containing OMP of capsular types A and D) or monovalent (containing OMP of P_52_ strain of *P. multocida*). The conventional alum adjuvanted bacterin vaccine prepared from P_52_ strain of *P. multocida* was used as the control vaccine. The outer membrane proteins of *P. multocida* have been studied by various workers for their importance in development of subunit vaccine as the outer membrane proteins are known to play important role in interaction of bacteria with the host’s epithelial cells and in virulence^4^. Though the outer membrane proteins are good immunogens, they require the help of an efficient adjuvant or delivery system that can aid in antigen uptake by antigen presenting cells. The alum based adjuvants have been used for many years but they face the drawback of causing site specific inflammatory reactions^24^. Nanoparticles based adjuvants are reported to efficiently stimulate the antigen uptake by antigen presenting cells^10, 26^.

The calcium phosphate nanoparticles were prepared using co-precipitation method. The maximum protein that could be loaded inside the calcium phosphate nanoparticle was 10 μg per mg of calcium phosphate nanoparticle-OMP conjugate. The nanoparticles were easy to disperse in aqueous medium. Joyappa *et al*. reported loading of 50μg plasmid DNA per mg of calcium phosphate nanoparticle-DNA conjugate. The variation of loading of DNA and protein (as in present study) could be due to the difference in biophysical properties of DNA and protein. The method of preparation of calcium phosphate nanoparticles is easier and can be efficiently used for production of vaccine in industrial scale. It takes nearly 18 hours to prepare the calcium phosphate nanoparticle-outer membrane protein conjugate. On the other hand, the double micro-emulsion method described by Bisht *et al*.^27^ for conjugation of calcium phosphate nanoparticle with DNA appears to be more difficult to perform and expensive.

The immunological studies indicated that the vaccine containing bivalent outer membrane proteins of capsular types A and D of *P. multocida* elicited better anti-capsular type A or D IgG response compared to that of the vaccine formulation containing outer membrane proteins of P_52_ strain. This could be due to significant antigenic variations observed in the outer membrane proteins of capsular types A, D or B of *P. multocida*. However, both the vaccine formulations were able to elicit 100%protection against both homologous or heterologous challenges. This could probably be due to the presence of common OMP band of relative molecular weight 47.95 kilodaltons in all the three capsular types. This protein might have some relationship with 47 kDa hypothetical protein reported by Wheeler (2009), which is known to be expressed by *P. multocida* in wide range of hosts^28^. It was reported to be one of the immunogenic porin proteins that is involved in long chain fatty acid transport across the cell membrane^25^. On the other hand, the bacterin vaccine containing alum adjuvant could not elicit significant level of protection against the challenges either with capsular types A or D of *P. multocida*. This could be due to masking of the immunogenic outer membrane proteins by the lipopolysaccharide of the outer membrane of the intact inactivated bacteria.

Although both the vaccine formulations containing either the OMP of P_52_ strain or bivalent OMP of capsular types A and D of *P. multocida* showed 100% protection, the later vaccine formulation showed significantly higher IgG response against capsular type A in mice.

As the first report of use of bivalent OMP based vaccine using calcium phosphate nanoparticles, the present study has showed that the vaccine formulation could elicit better immune response compared to that of alum adjuvanted bacterin in mice. Further study is required to assess its protective efficacy in pigs against swine pasteurellosis.

## ACKNOWLEDGEMENT

The authors are grateful to the Centre Director, DBT-AAU Centre, Assam Agricultural University, Jorhat, Assam, India for providing financial support to carry out the work. The authors are also grateful to the Advanced State Biotech Hub, College of Veterinary Science, AAU, Guwahati for providing the laboratory facilities.

## Notes

### Competing Interest Statement

The authors have declared no competing interest.

